# A time-resolved imaging-based CRISPRi screening method

**DOI:** 10.1101/747758

**Authors:** Daniel Camsund, Michael J. Lawson, Jimmy Larsson, Daniel Jones, Spartak Zikrin, David Fange, Johan Elf

## Abstract

Our ability to connect genotypic variation to biologically important phenotypes has been seriously limited by the gap between live cell microscopy and library-scale genomic engineering. Specifically, this has restricted studies of intracellular dynamics to one strain at a time and thus, generally, to the impact of genes with known function. Here we show how *in situ* genotyping of a library of *E. coli* strains after time-lapse imaging in a microfluidic device overcomes this problem. We determine how 235 different CRISPR interference (CRISPRi) knockdowns impact the coordination of the replication and division cycles of *E. coli* by monitoring the location of replication forks throughout on average >500 cell cycles per knockdown. The single-cell time-resolved assay allows us to determine the distribution of single-cell growth rates, cell division sizes, and replication initiation volumes. Subsequent *in situ* genotyping allows us to map each phenotype distribution to a specific genetic perturbation in order to determine which genes are important for cell cycle control. The technology presented in this study enables genome-scale screens of virtually all live-cell microscopy assays and, therefore, constitutes a qualitatively new approach to cellular biophysics.

## Introduction

The last decade has shown remarkable development in genome engineering, fronted by applications of Cas9-mediated gene targeting ^1, 2^. In combination with inexpensive large-scale DNA oligonucleotide synthesis, these techniques make it possible to generate pool-synthesized cell strain libraries with specific perturbations genome-wide ^3, 4^. More recently, methods have been developed for screening Cas9 genome edited libraries by sorting the library members based on the expression of a fluorescent protein and then sequencing cells with similar expression level ^5^ or using single-cell RNAseq on libraries of CRISPR interference (CRISPRi) perturbations to determine the state of the transcriptome for individual cells ^6–8^. These methods are however blind to cellular dynamics and intracellular localization of relevant molecules.

The progress in genome-scale engineering and expression perturbation has been accompanied by equally impressive developments in microscopy and microfabrication, which enable characterization of complex phenotypes at high temporal and spatial resolution in living cells under well-controlled conditions ^9–15^. While the power of these methods enables deep insight into cellular biophysics, the limitation of working with one strain at a time prohibits studying the impact of genes whose function is not already, at least to some degree, known. Given the rapid progress within the previously separate fields of imaging and genomics, the lack of efficient techniques for time-resolved single-cell phenotyping of pool-synthesized genetic strain libraries constitutes a severe bottleneck in biological research. We have recently proposed a tentative solution to the problem ^16^ that we now demonstrate scaled to hundreds of genes for tens of thousands of single cells, converting the concept into a practical tool. We use the method to perform a large-scale screen of complex phenotypes to identify the regulatory elements of replication-division coordination in bacterial cells.

## Results

### Overview of the method

The heart of our method is a microfluidic device that enables both high-resolution dynamic phenotyping and subsequent *in situ* genotyping of the individual strains. The microfluidics approach allows us to keep the bacteria in a constant state of exponential growth over hundreds of generations while imaging them at high resolution. The fluidic device ^13^ is an adaption of the “mother machine chip” ^15^ where each cell trap has a 300 nm constriction at the end that enables fast loading, media and probe exchange (see the left side of Fig. 1 for a schematic of the chip). Each strain occupies a defined space in the fluidic chip, but the genotypes of the different strains are unknown at the time of phenotyping. After the phenotypes have been determined, the cells are chemically fixed in the chip, and the genotypes are optically inferred by sequential fluorescence *in situ* hybridization (FISH) to a barcode (Fig. 1). We refer to the technique as DuMPLING (Dynamic u-fluidic Microscopy Phenotyping of a Library before IN situ Genotyping).

**Figure 1.**
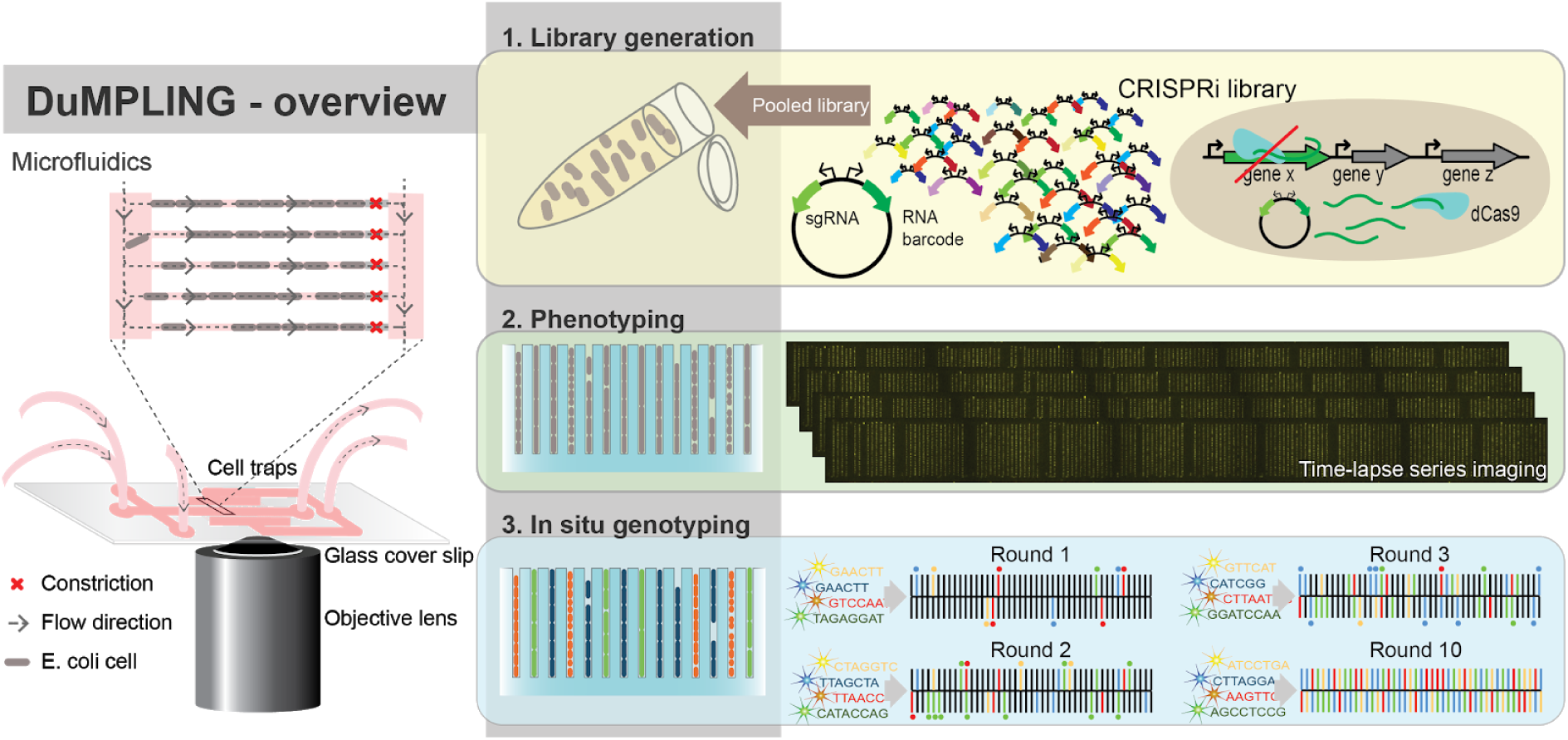
Assay workflow. **1.** The bacteria in the CRISPRi library contain pool-synthesized plasmids each expressing a barcode and a corresponding sgRNA (top) for repressing a specific gene in the *E. coli* chromosome. The library of cells is loaded into a fluidic device where each strain occupies a spatially separated trap (left) and where **2.** the cells can be monitored with highly sensitive time-lapse fluorescence microscopy for hours or days. **3**. After phenotyping the cells are fixed, and the identities of the strains are revealed by sequential FISH probing for the barcodes.

We used the DuMPLING technique to characterize the coordination of the replication and division cycle in *E. coli* by tracking the chromosome replication forks throughout the cell cycle in a CRISPRi library. Replication initiation in *E. coli* is triggered at a fixed volume per chromosome ^17, 18^ independent of growth rate, but the underlying molecular mechanism is largely unknown. In this work, replication initiation was studied directly by observing a strain with a chromosomally integrated *seqA-yfp* fusion. SeqA binds hemimethylated DNA in the wake of the replication machinery and can thus be used to track replication foci (Fig. 2). In addition, the cell size at division and replication initiation was determined using phase contrast imaging. By imaging hundreds of cell cycles for each CRISPRi perturbation, we aimed to identify which genes are involved in setting the accuracy of the initiation volume. This screen could not have been performed without monitoring the dynamics of replication initiation directly in individual cells.

**Figure 2.**
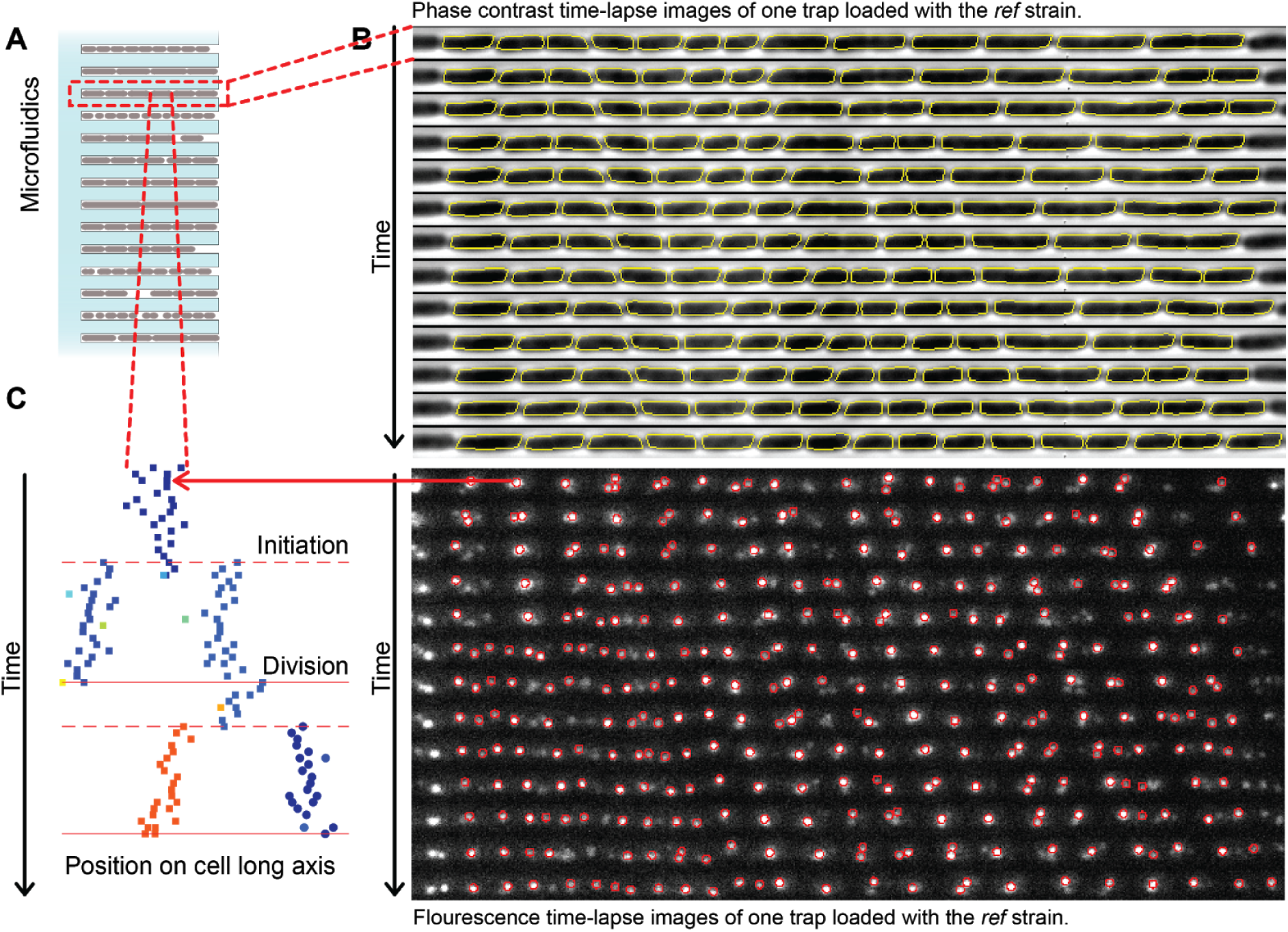
Analysis. **A.** Cartoon of cells growing in a microfluidic chip. **B.** Example kymographs in phase contrast (top) and fluorescence (bottom), with automated cell segmentation (*yellow*) or detection of SeqA-YFP clusters (*red circles*), respectively, overlayed. **C.** The time series of cell segmentation and SeqA-YFP clusters are combined to generate single cell trajectories across cell divisions and chromosome replications.

### Implementation and Analysis

We constructed the library in a host strain of MG1655 with a *seqA-yfp* fusion for tracking the replication forks, dCas9 expressed from an anhydrotetracycline (aTc) inducible promoter for CRISPRi, and a T7 RNA polymerase (T7 RNApol) expressed from an arabinose-inducible promoter for barcode RNA expression. Each of the 235 library members expressed a unique single guide RNA (sgRNA), directing dCas9 to bind and repress a specific gene (Online Methods 1.1.2). The library included all known, non-lethal cell cycle-related targets as well as 38 y-genes ^19^, 28 of which are largely uncharacterized or have an unknown function. By inducing the dCas9 expression with 1 pg/ul aTc, the target genes were downregulated between 10-100 times (Supplementary Text 2.2). Each plasmid also encoded a 20 bp-barcode sequence that could be expressed as RNA from an inducible T7 promoter. The barcode was uniquely coordinated with the sgRNA (Online Methods 1.1.4) sequence to identify which sgRNA was expressed in which strain.

We loaded 40 pool synthesized strains at a time into the microfluidic device in order to phenotype on average 562 generations per strain in an 8 h experiment. Before phenotyping, the bacteria were grown with dCas9 induced for 9 hours in order to establish steady state phenotypes. This time also allowed the mother cell at the end of the cell trap to divide enough times to set the genotype of the whole trap. To achieve sufficient time resolution, we imaged each position (30 traps per position) for 20 ms every minute in the phase contrast channel and 300 ms every two minutes in the fluorescence channel (514 nm@5.3 W/cm^2^). This limited us to 90 positions, or 2700 cell traps per experiment. This also gives enough traps per strain to account for uneven representation of individual genotypes in the library (Fig. S1). The limitation for how many strains we can analyze simultaneously is currently set by the number of traps we can image during phenotyping (e.g. how fast we can image and move the stage) and not by how many strains can be made or genotyped in parallel.

After phenotyping, the barcode RNAs were expressed by T7 RNApol induction and the cells were subsequently fixed and permeabilized. The strains were identified *in situ* by sequential FISH probing in four colors; each round of probing identifies the positions of four unique strains. Each round was completed in less than 30 mins, since no stripping of probes was required, and all strains could be identified within 6 hours following fixation. In complementary experiments where we loaded all strains simultaneously, we used combinatorial FISH probing which can identify N^R^ genotypes in R rounds of N colors, but this approach is more time-consuming at this library-scale since the probe stripping is slow and the readout probe concentrations are lower (see Supplementary Text 2.3 for additional information about advantages and drawbacks of different genotyping methods in the PDMS chip).

After connecting the genotypes to the phenotypes through their spatial position, we analyzed how the repression of individual genes affected the cell size and growth rate (Fig 2B top) as well as coordination of replication with the division cycle (Fig 2B bottom). The kymograph for one of the 21700 analysed cell traps is displayed in Fig 2C. The bacteria were segmented using the POE method ^20^ and tracked by the Baxter algorithm ^21^. The SeqA-YFP foci were detected by a wavelet based method ^22^ and the replication forks were tracked simultaneously through the generations using the u-track algorithm ^23^ (Fig 2C). Each experiment, including 2700 cell traps that were imaged every minute for 8 hours, resulted in 220 Gb of image data and took 2 hours to analyze on 45 cores using customized parallelized image analysis routines.

### The impact of perturbations on the *E. coli* cell cycle

In Fig 3A-C we show comparisons of the average growth rates and cell sizes at division and replication initiation respectively for the 215 strains for which we obtain data from a minimum of 5 independent cell traps and 40 complete cell cycles. The genotypes that are substantially different from the reference control strain in replicate experiments (*ref*, see Online Methods 1.1.4 for strain details) are indicated by the name of the sgRNA targeted gene. The sizes at division and initiation can get both bigger and smaller than *ref*, whereas growth rates typically only get smaller. The deviations from *ref* are mostly uncorrelated between the properties, with two notable exceptions: the *tol-pal* cluster is smaller and the *fis, diaA*-cluster is larger in both initiation and birth volumes (Fig. 3A).

**Figure 3.**
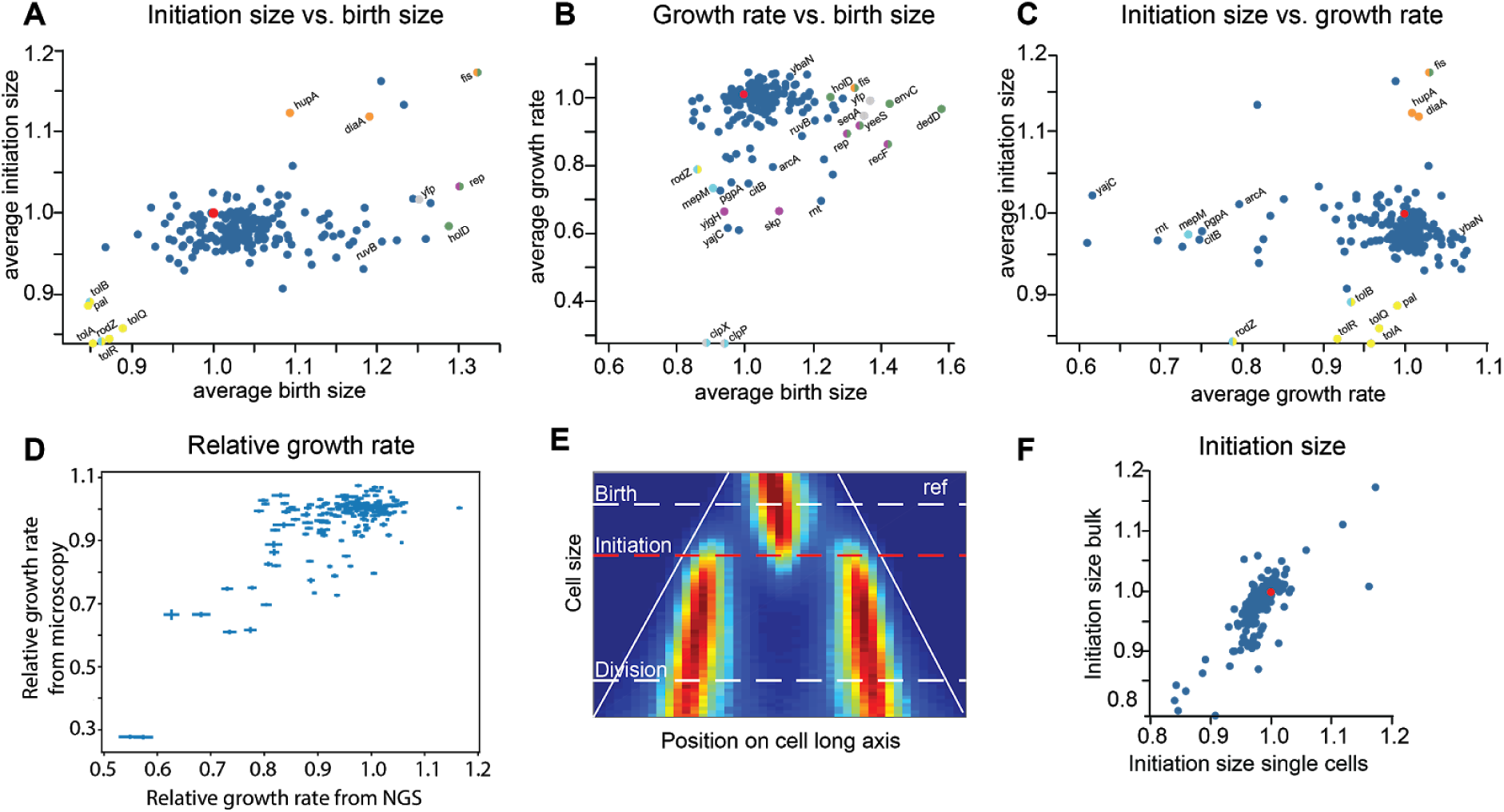
Phenotypic data averaged for each genotype. **A-C** Two dimensional plots in phenotype space, CRISPRi knockdowns with significant deviation from the reference control strain (*ref*) are labeled by the name of the targeted gene. Outlier dots have been classified and labelled as follows: large birth size (green), small birth size (cyan), large initiation size (orange), small initiation size (yellow), no foci (grey) and undefinable initiation size (purple). As these properties are not all mutually exclusive, dots may by multicolored. **A.** Horizontal: Average normalized birth size. Vertical: Average normalized initiation size. **B.** Horizontal: Average normalized birth size. Vertical: Average normalized growth rate. **C.** Horizontal: Average normalized growth rate. Vertical: Average normalized initiation size. **D.** Control experiment showing that the relative growth rate derived from the DuMPLING experiments correlate well with that from a pooled competition assay (correlation = 0.7). **E.** Fork distribution plot. Horizontal is SeqA-YFP cluster location along the long axis of the cell(from old cell pole to new), vertical is cell size, color indicates the probability of finding a replication fork at a given position along the cell axis and a given cell size. Initiation size corresponds to the average of individually tracked replication forks. **F.** Average normalized initiation size fit to bulk replication forks (vertical) compared with the average of single cell initiation size estimates (correlation = 0.76).

As a control for correct genotyping and cell segmentation, the average growth rates obtained from the single-cell time-lapse imaging compare well to the corresponding bulk experiment (Fig. 3D). To make bulk estimates of the growth rates, we performed a competition assay of the whole library in liquid culture and determined the time-dependent relative abundance of each genotype by next generation sequencing (NGS) (Fig 3D, assay details in Supplementary Text 2.1). Additional controls are described in the Supplementary Text 2.4.1-2.4.6. For example, we compared the phenotypes of selected strains from the DuMPLING screen to those obtained with the corresponding knock-outs or specific sgRNA knock-downs constructed and measured one strain at a time (Supplemental Text 2.4.4). The good agreement in phenotypes implies that the genotyping is robust, although there were notable exceptions with off-target effects or where synthesis errors in the sgRNA coding sequence were selected for.

An information-rich way to simultaneously characterize replication and cell cycle processes in individual strains is the fork distribution plot, which shows the probability of finding a replication fork in a specific position in the cell (horizontal axis) for cells of a given size (vertical axis). The distribution for the *ref* strain is shown in Fig 3E. The data in the fork distribution plots can be used to estimate average initiation volume (see Fig. 3E, Online Methods 1.3.1 and ^17^). The good agreement between this bulk estimate and the average of estimates obtained from individual cells (see Fig. 3F for comparison and Online Methods 1.3.1 for estimation of a single cell initiation size) acts as a control for our ability to accurately estimate initiation size for individual cell trajectories.

Fig. 4A show the fork distribution plots for all genotypes from one experiment that met the criteria described in the figure legend. Most of these are similar to the unperturbed *ref* (Fig 3E), but a number of variant classes can be identified: Fig 4B (*fis, dedD* etc.) represents strains with large division size and Fig 4C (*clpP, nfh, pal* etc.) strains with small division size. Correspondingly Fig 4D (*fis, ihf* etc.) represents strains with large initiation size and Fig 4E (*tol, pal* etc.) represents strains with small initiation size. The strains in Fig 4F (*seqA, dam* and *damX*) have no SeqA-YFP foci, which makes sense since *dam* encodes the DNA methylase and SeqA only binds hemimethylated DNA. *damX* is located upstream of dam in the same operon and its repression will also downregulate *dam*. Fig 4G (*hda, ihf* etc.) represents strains for which the replication initiation size is undefined, suggesting that there may be significant cell-to-cell variability in the initiation size and consequently, that these genes are important for regulation of DNA replication.

**Figure 4.**
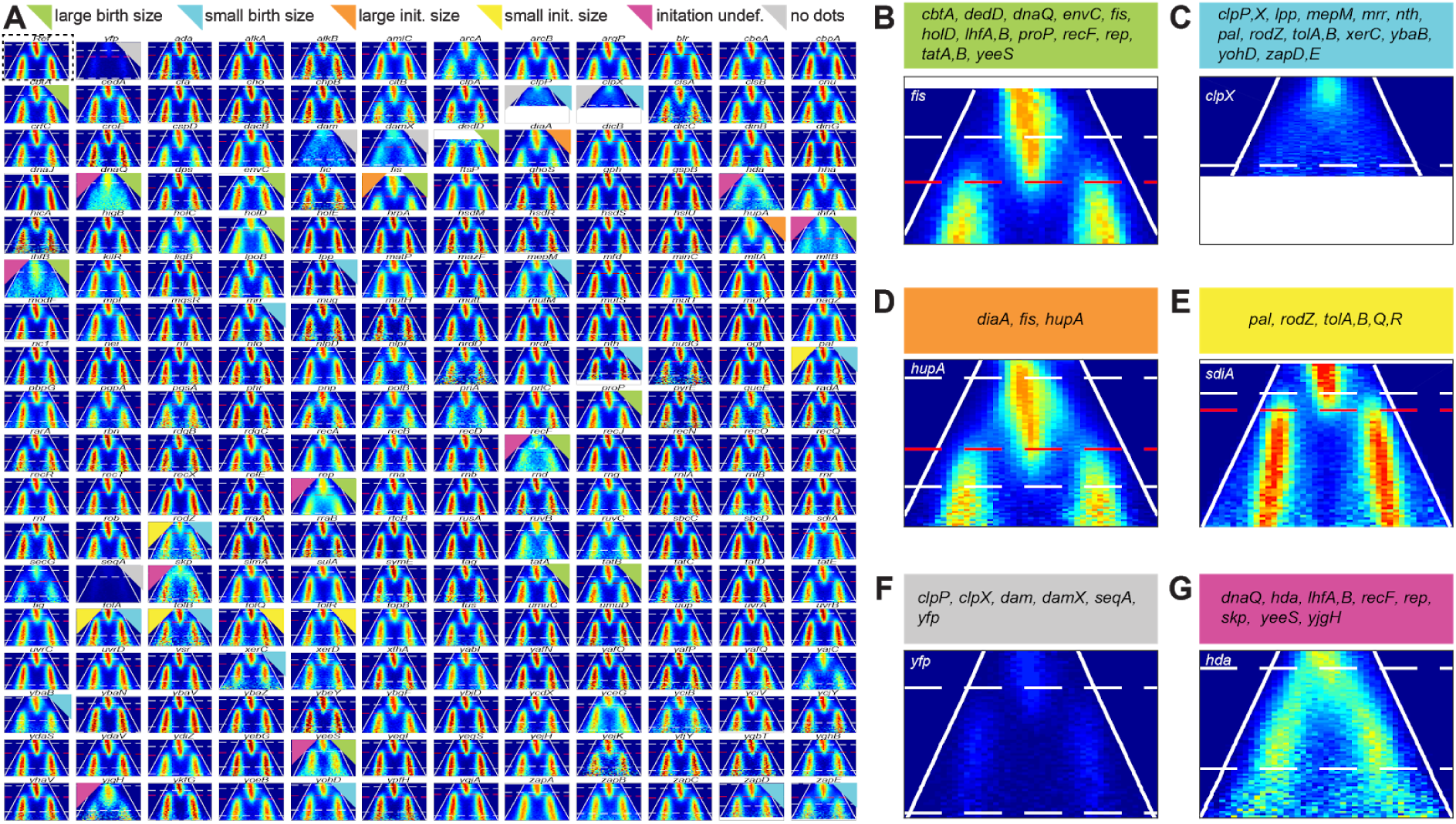
Fork distribution plots and time-resolved phenotypes. **A.** Plots for all knockdowns that had a minimum of 5 independent cell traps and 40 whole cell cycles. Color labeling indicates the classification of phenotypes. Fork distribution plots are laid out as in Fig. 3E. *White dashed lines*: Birth/division size. *Red dashed lines*: replication initiation size, average of individually tracked replication forks. **B-G:** Examples of each classification from **A** (NB: it is possible for a gene to belong to more than one classification). **B.** Large division size (only birth size in figure). **C.** Small division size (only division size in figure). **D.** Large initiation size. **E.** Small initiation size. **F.** No SeqA-YFP foci. **G.** Replication initiation size not definable. Fork distribution plots for a replica experiment is shown in figures S2-S7.

By pooling data from different cells of the same genotype, we lose information about cell-to-cell variation for the different genotypes. As a first-order description of the cell-to-cell variation, we plotted the coefficient of variation (CV) against the corresponding average in growth rate, birth size and initiation size for each genotype in Fig 5A-C. Corresponding illustrative examples of the full distributions are also given for selected genotypes in Fig 5D-F. For birth size most perturbations give larger average as well as CV (Fig. 5B). Interestingly *fis* and *dedD* do however increase the average size without an increase in CV. In a few cases the CV increases much more than the change in the average. For example, *hda* repression gives a 80% increase in variation of the replication initiation size with only a 10% reduction in the average (Fig. 5C), suggesting an important regulatory role. With respect to cell-to-cell variation in growth rate, we observe a striking trend that gene knockdowns that reduce growth rate also give rise to large cell-to-cell variation (Fig. 5A). A plausible explanation is that the repression makes cell growth limited in only one factor and that stochastic fluctuations in this factor alone directly impact the growth rate.

**Figure 5.**
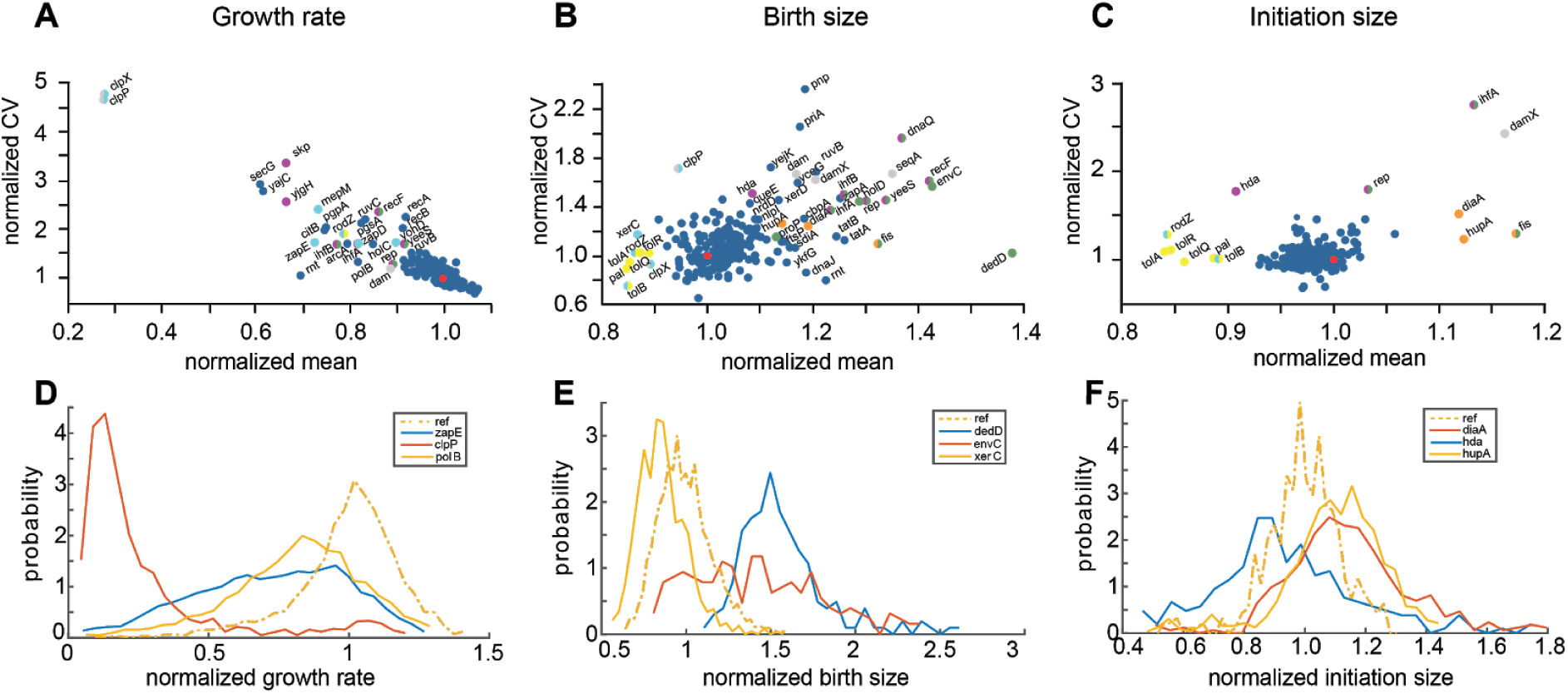
Cell-to-cell phenotypic variation for each genotype. *Top row*: Each plot has the coefficient of variation (CV) on the vertical axis and mean on the horizontal axis, both normalized by *ref* (red dot at [1,1] in each plot), for **A.** growth rate, **B.** birth size and **C.** initiation size. *Bottom row*: In D-F we show probability density (vertical axis) as a function of **D.** growth rate, **E.** birth size and **F.** initiation size for selected genotypes.

## Discussion

Circling back to the original question of which genes are important for triggering replication at a fixed cell size, a number of candidates stand out in Fig 4G and 5C. If we exclude genes with well documented and clearly unrelated functions, we end up with *hda, diaA, ihfA, ihfB, fis*, *yjgH* and *yeeS. diaA* is previously known for its direct interaction with DnaA at replication initiation ^24^. *hda, ihfA, ihfB,* and *fis* are all associated with DnaA-ATP to ADP cycling ^25^. Further characterization of *yjgH* shows that the observed phenotype is due to an off-target effect of the sgRNA (Supplementary Text 2.4.3). The remaining uncharacterized gene that shows an evident perturbation in the accuracy of replication is *yeeS*. This gene deserves more specific study, although there is a chance that the perturbation is mediated by the CbtA toxin ^26^ that is encoded on the same transcript as *yeeS.* We also note that downregulation of *pgsA* does not show a perturbation in replication initiation although the gene product is key in the synthesis of fatty acids, which some studies have implicated in DnaA-ADP to ATP conversion at the membrane ^27^. Thus, overall, our assay supports models for replication initiation control based on DnaA-ATP to ADP cycling through the regulatory inactivation of DnaA (RIDA) mechanism mediated by Hda ^28^ as well as at the DnaA-reactivating sequences (DARS) ^29^ binding Fis and Ihf and *datA* locus dependent DnaA-ATP hydrolysis (DDAH) ^30^ also regulated by Ihf.

The DuMPLING approach, here deployed to identify the key regulators of the *E. coli* cell cycle, can be used to study all sorts of complex dynamic phenotypic traits that require sensitive or time-lapse microscopy for characterization. Plausible extensions include studies of gene expression dynamics in response to recoded promoter sequences or other genetic regulatory elements; live cell enzymatic assays as a function of active site residue mutations; as well as interaction partner screens that alter the intracellular location of a labeled reporter molecule in response to all possible gene knockdowns.

## Acknowledgments

This work was supported by the Knut and Alice Wallenberg foundation (KAW) and the Swedish research council (VR). We are grateful to Dr. Irmeli Barkefors for help with the manuscript and figures and to Praneeth Karempudi for making microfluidic molds.

## Author contributions

JE conceived the DuMPLING method and SeqA application, DC developed cloning methods, designed and made strain library, JE and ML managed the project, DF and ML developed phenotyping methods, JL and ML developed genotyping methods, DF built the microscope, JL and ML made microscopy experiments, SZ, DF developed image analysis pipeline, SZ, DF analysed DuMPLING data, DJ made repression measurements and NGS growth rate experiments, JE, ML, DC and DF wrote the paper with input from all authors.

## Data and code

All raw data and analysis codes will be publically deposited at the time of publication.

